# Inference of cell-cell interactions from population density characteristics and cell trajectories on static and growing domains

**DOI:** 10.1101/011080

**Authors:** Robert J.H. Ross, C.A. Yates, R.E. Baker

## Abstract

A key feature of cell migration is how cell movement is affected by cell-cell interactions. Furthermore, many cell migratory processes such as neural crest stem cell migration [1, 2] occur on growing domains or in the presence of a chemoattractant. Therefore, it is important to study interactions between migrating cells in the context of domain growth and directed motility. Here we compare discrete and continuum models describing the spatial and temporal evolution of a cell population for different types of cell-cell interactions on static and growing domains. We suggest that cell-cell interactions can be inferred from population density characteristics in the presence of motility bias, and these population density characteristics for different cell-cell interactions are conserved on both static and growing domains. We also study the expected displacement of a tagged cell, and show that different types of cell-cell interactions can give rise to cell trajectories with different characteristics. These characteristics are conserved in the presence of domain growth, however, they are diminished in the presence of motility bias. Our results are relevant for researchers who study the existence and role of cell-cell interactions in biological systems, so far as we suggest that different types of cell-cell interactions could be identified from cell density and trajectory data.

## 1 Introduction

It is widely understood that cell-cell interactions play an important role in cell migration [3–6]. For example, multiple different cell-cell interaction mechanisms have been identified as promoting metastasis in breast cancer [3], and repulsive interactions mediated via ephrins on the surface of neural crest stem cells (a cell population that play a fundamental role in vertebrate development) have been shown to be important in organising neural crest stem cell migration [4]. Therefore, understanding interactions between motile cells is essential for understanding cell migration in biological systems [7,8]. However, although great progress has been made in the identification and quantification of cell-cell interactions, work to understand the effect different cell interactions have on cell motility at both individual and population levels is still ongoing [9–14]. Mathematical and computational modelling allows us to assay a large range of cell-cell interactions with complete control over parameters, and perform simulations in environments that are difficult to reproduce or control experimentally, such as domain growth. This allows us to test multiple hypotheses with relative ease.

Throughout this work we use mathematical and computational modelling to test simple measurements that could be carried out during experiments in order to discern cell-cell interactions. To do so we use a cell-based discrete random-walk model with volume exclusion to represent a two-dimensional motile cell population [15]. Discrete random walk models based on a simple exclusion process have been widely applied to the study of biological phenomena [16–20]. For instance, discrete random walk models have been used to study spatial structure in cell aggregations [16, 17], the growth of cell colonies [18], enteric colonic growth [19], infection spreading and ecological interactions [20]. We use this model as it is straightforward to incorporate a large number of cell-cell interactions, the focus of our study [3, 4, 12, 21–23]. We also study these cell-cell interactions in the context of growth and motility bias. Motility bias represents the directed migration of a cell caused, for example, by the presence of a chemoattractant [24–26]. Chemoattractants have been shown to play a major role in cell migration. For instance, chemoattractants are known to be essential for the correct migration of cranial, cardiac and enteric neural crest stem cells [1, 2, 25, 27]. In addition, growth is an essential component of biological systems, especially in the context of development, when large scale cell rearrangements are taking place [28]. This means that the inclusion of domain growth into models is necessary to understand cell migration in many contexts.

Importantly, we derive continuum equivalents of our discrete models for the spatial and temporal evolution of the cell population density, and the expected displacement of tagged cells on both growing and static domains [21–23, 29]. These continuum approximations in many cases are extensions of previously derived continuum models [21–23]. Their derivation is important because discrete models are often computationally expensive and analytically intractable. This means that explaining discrete model behaviours and parameterising them using e.g. Monte Carlo methods [30, 31] is difficult.

By implementing simple measurements on our simulation results we show that cell-cell interactions can be inferred from population density characteristics in the presence of motility bias, and these population density characteristics for different cell-cell interactions are conserved on both static and growing domains. Similarly, different cell-cell interactions can be inferred from the characteristics of the trajectory of a tagged cell. The results presented here are relevant to experimentalists in the field of cell biology, as the study of population density and cell trajectory characteristics could be used to distinguish simply between different cell behaviours and interactions.

The outline of this work is as follows. To begin we introduce a generic discrete model for cells migrating on a growing domain. We also define mathematically different types of cell-cell interactions for our discrete model, and derive corresponding continuum approximations of our discrete models. We then compare the accuracy of these continuum approximations with their discrete model counterparts. We do this for a range of cell-cell interactions, for both static and growing domains. Finally, we show that different cell-cell interactions can be inferred from population density and cell trajectory characteristics using simple measurements.

## 2 Model formulation

In this section we first introduce our discrete modelling framework and the cell-cell interactions that we employ throughout this work. We then derive continuum approximations of our discrete models, and compare the discrete model simulation results with our continuum approximation solutions.

### 2.1 Discrete modelling framework

We use an agent-based, discrete random-walk model on a two-dimensional square lattice with lattice spacing Δ [20] and size *L*_*x*_*(t)* by *L*_*y*_*(t)*. For discrete simulations in which the domain does not grow, the lengths of the *x* and *y* axes are constant, *L*_*x*_*(t)* = *L*_*x*_ and *L*_*y*_*(t)* = *L*_*y*_. All simulations are performed with periodic boundary conditions at *y* = 0 and *y* = *L*_*y*_*(t)*, and no-flux boundary conditions at *x* = 0 and *x* = *L*_*x*_*(t)*. Furthermore, in all simulations displayed here, domain growth only occurs along the *x* axis. However, the growth mechanism presented here is equally applicable in two dimensions [19].

The simulation runs with a fixed time step of duration *τ*. The lattice is populated by agents, which represent biological cells. Each agent is assigned to a lattice site, from which it can move into an adjacent site. If an agent attempts to move into a site that is already occupied, the movement event is aborted. This process, whereby only one agent is allowed per site, is typically referred to as an exclusion process [20]. In this work we neglect cell proliferation, death and differentiation as they add significant complications to our methodology and provide little extra insight.

At each time step, all cells on the lattice have the opportunity to move to an adjacent site with probability *P*. On a square lattice, with a cell positioned at site (*row* = *i*, *column* = *j*) and not on a boundary, adjacent sites would consist of sites (*i, j* − 1), (*i* − 1, *j*), (*i, j* + 1) and (*i* + 1, *j*). To relate site coordinates to Cartesian co-ordinates, we use *x* = *i*Δ, *y* = *j*Δ. This means the length of the *x* axis is *L*_*x*_(*t*) = *N*_*x*_(*t*)Δ, where *N*_*x*_(*t*) is the number of elements in the *x* axis. For all simulations presented here *P* = 1. The random sequential update method we employ means that if there are *M* cells in the domain, *M* sequential random cell movements to an adjacent site are attempted at each time step with replacement [15]. This means each cell is chosen on average once per time step, however, in some time steps a cell may be selected more than once or not at all.

Key to both the discrete and continuum models we derive throughout the course of this work is the growth mechanism we employ. This growth mechanism, which we shall refer to as a ‘pushing’ mechanism, has been successfully used in modelling avian gut growth [19], and to compare the accuracy of different stochastic update schemes for discrete models [29, 32]. Our pushing growth mechanism is as follows: at each time step one site is randomly selected from each row of the domain to undergo growth. This means that all rows remain the same length throughout the simulation, and that domain growth is linear. It also means that all discrete realisations grow to the same length, and so in this respect our growth mechanism is deterministic. We choose linear domain growth as it has been shown to be present in many biological systems [2, 19, 33, 34]. However, this growth mechanism is equally applicable to other types of isotropic domain growth, such as exponential growth [29]. If site (*i*, *j*) is selected for growth, the new site is inserted at (*i*,*j*), and the ‘selected’ site is moved to (*i*, *j* + 1), taking its contents (i.e. an agent or no agent) with it. This means new sites are unoccupied initially, and that all sites to the right of a site that undergoes growth are ‘pushed’ one lattice spacing, Δ, in the positive *x* direction, along with their contents. This growth mechanism is illustrated in Fig. 1.

**Figure 1:**
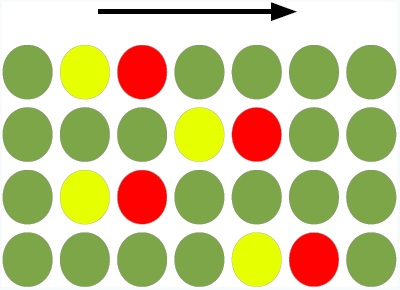
(Colour online). ‘Pushing’ growth mechanism. Growth in the *x* axis for a two dimensional lattice. The arrow indicates the direction of growth. In each row the red (dark grey) site has been chosen to undergo a growth event, whereby the red site moves one lattice spacing to right, carrying its contents. Consequently, all sites to the right of a red (dark grey) site move one space to the right as well. The new site (yellow, light grey) is inserted where the red site was at the start of the time step. This new site (yellow, light grey) is unoccupied.

### 2.2 Transition probabilities that represent cell-cell interactions

As outlined in the introduction it is known that cell-cell interactions play an important role in cell migration [3, 5, 6]. Therefore, we implement a number of cell-cell interactions in both our discrete and continuum models. These four types of cell-cell interaction, blind, adhesive, repulsive and myopic, imitate cell-cell interactions in biological systems [3, 5, 22, 22, 23]. We also study the effect of motility bias, as it is known to play a crucial role in cell migration [24, 26, 27]. It is important to note that the way in which we model these cell-cell interactions is not unique. However, given the focus of our study to discriminate between different types of cell-cell interactions we have chosen models of cell-cell interactions that have previously been published, and in some senses can be seen as ‘canonical’ examples.

To define our cell-cell interactions we first introduce the notation we will use [23]. For any site *v* on a square lattice we define the nearest neighbourhood set 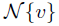. This set contains all lattice sites that share a boundary with *v*, that is 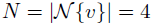 (away from the boundary) for a two-dimensional square lattice. This neighbourhood is typically referred to as a von Neumann neighbourhood, and these are the sites from which agent movements into and out of site *v* can be made.

Throughout this work we use standard conservation arguments to derive our continuum approximations [35, 36]. The occupancy of site *v* is denoted by *C*_*v*_, with *C*_*v*_ = 1 for an occupied site and *C*_*v*_ = 0 for an unoccupied site. By averaging over many identically prepared realisations of our discrete system we obtain the expected occupancy of site *v*, denoted by 〈*C*_*v*_〉. This quantity represents the probability of occupancy of site *v*. We will use these expected occupancies to derive continuous approximations of our discrete models. Importantly, we assume that adjacent sites are independent in derivation of the continuum approximations. This approach is well-established in the literature [21, 23, 37, 38].

#### 2.2.1 Blind agent interactions

In ‘blind’ agent interactions the agent does not consider the occupancy status of adjacent sites before attempting a move. In this sense the agent is ‘blind’ to its environment. The transition probability for a blind agent, in which an agent attempts to move to one of the *N* nearest neighbour sites with probability *P*(1 + *ρ*_*v→v′*_)/*N* is written as

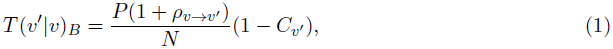

such that

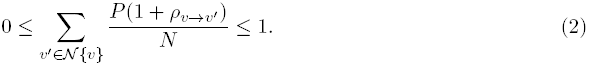

In Eq. (1) *T*(*v′*|*v*)_*B*_ is the probability that an agent will successfully move from site *v* to site 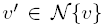 during [*t*, *t* + *τ*), given it is at *v* at time *t* (the subscript *B* denotes blind) [21]. *ρ*_*v→v′*_ is the bias present in moving from site *v* to *v′*, *ρ*_*v→v′*_ ∈ [-1, 1] and Eq. (2) constrains the values *ρ*_*v→v′*_ can take. That is, the bias terms (and *P*) must be chosen so that Eq. (2) is satisfied. For our purposes *ρ*_*v→v′*_ will be one of *±ρ*_*x*_ or *±ρ*_*y*_, with *±ρ*_*x*_ the bias for movements in the positive/negative *x* directions and *±ρ*_*y*_ for movements in the *y* direction. For example, a movement from site (*i, j*) →(*i, j* + 1) would have a bias of *ρ*_*x*_. Conversely, a movement from site (*i, j*) → (*i, j* − 1) would have a bias of −*ρ*_*x*_.

We now proceed to define more complicated agent interactions. To do so we adopt a transition probability notation similar to that used by Landman and Fernando [23], with some modifications. From our nearest neighbourhood set 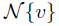 we define two further sets. 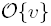 is defined as the set of all nearest neighbour occupancy configurations in the set 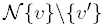. 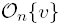 is then defined as a subset of 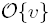 containing exactly *n* occupied nearest neighbour sites from the possible *N* − 1 sites in the set 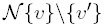. We then use 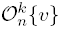 to index the possible choices of sites in 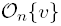, where the superscript *k* denotes one of the 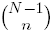 possible choices of *n* distinct sites in 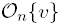. Put simply, these represent all the possible combinations of occupied sites in 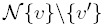 and are theelements of set 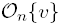. Additionally, we define 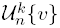 to be 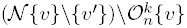. Therefore, 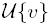 can be understood as the set of all possible combinations for unoccupied sites in the nearest neighbourhood set of site *v*. These combinations are displayed in Fig. 2. With this notation we can now define more complicated agent interactions for our discrete model.

**Figure 2:**
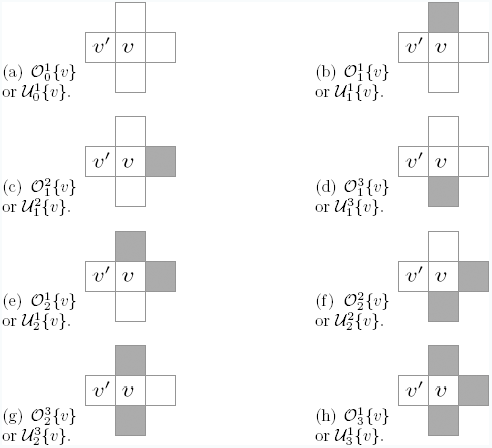
The nearest neighbourhood occupancy sets. The shaded sites indicate a site that is occupied by a cell.

#### 2.2.2 Adhesive/repulsive agent interactions

In adhesive and repulsive agent interactions, the number of sites that are occupied in the nearest neighbourhood of an agent influence the probability of that agent completing a move. In this sense the agent is not ‘blind’, as it takes its nearest neighbourhood into account before attempting a move. For adhesive reactions the success of a move decreases if the agent is in contact with other agents that it can adhere to. Conversely, for repulsive interactions the success of a move increases if an agent is in contact with other agents. In adhesive and repulsive agent interactions an agent attempts to move to one of its *N* nearest neighbours with probability of *P*(1 + *ρ*_*v→v′*_(1+α)*^n^*/*N*, where *n* ≤ *N* signifies the number of nearest neighbours of *v* that are occupied, −1 ≤ *α* < 0 for adhesive agent interactions, and 0 < *α* ≤ 1 for repulsive agent interactions. This transition probability was first suggested by Khain et al. [22], and has been successfully applied to discrete models of wound healing and cell aggregation [22, 36, 39].

We denote the occupancy of sites *u* and *w* by *C*_*u*_ and *C*_*w*_, respectively, and let 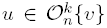 and 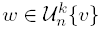 represent all configurations of occupied and unoccupied members of the nearest neighbour set of site *v*, respectively. For a two-dimensional lattice the transition probability for adhesive/repulsive agent interactions with bias can be written as

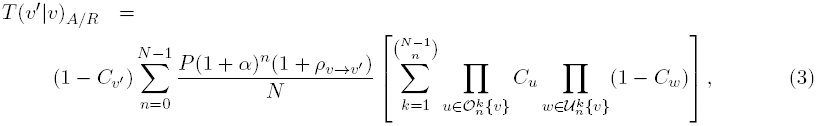

such that

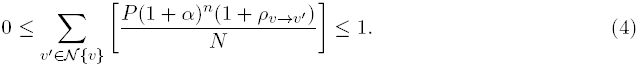

Here, the subscript *A/R* denotes adhesive/repulsive agent interactions (adhesion or repulsion depends on the sign of *α*). Note that if *α* is zero Eq. (3) is equivalent to Eq. (1), the blind transition probability. This is because the sum of the probabilities of all possible neighbourhood configurations is equal to unity. Once again, condition Eq. (4) constrains the values *P*, *α* and *ρ*_*v→v′*_ can take in Eq. (3).

#### 2.2.3 Myopic agent interactions

The final interaction type we study is myopic agent interactions. A myopic agent never attempts to move into an occupied site [40]. Instead, an agent’s probability of moving to an occupied site is redistributed amongst non-occupied sites. That is

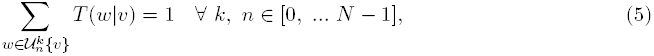

for an agent selected to move in the discrete model. For a transition probability describing myopic agent interactions without bias this means an agent at site *v* attempts to move into an unoccupied site in 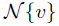 with equal probability *P*/(*N* − *n*), where *n* signifies the number of nearest neighbours of *v* that are occupied. If bias is introduced, then not only is the number of occupied neighbours important, but also the position of the occupied neighbours. An example of myopic agent interactions is displayed in Fig. 3. For Fig. 3 (a), none of the nearest neighbour sites of *v* are occupied, so an agent at *v* would move to site *v′* with probability

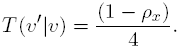

However, in Fig. 3 (b), a site in the nearest neighbourhood of *v* is occupied, and so the probability of moving to this occupied site from *v* must be redistributed amongst the unoccupied sites. Therefore, an agent at site *v* would move to site *v′* with probability

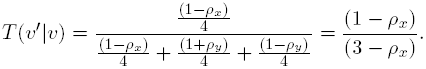

**Figure 3:**
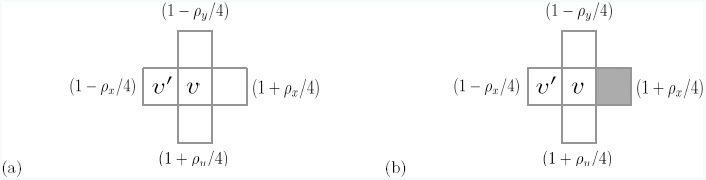
Myopic neighbourhood example. In panel (a) an empty nearest neighbourhood is displayed. With this configuration *T*(*v′*|*v*) = (1−*ρ*_*x*_)/4. In panel (b), a nearest neighbourhood in which one site is occupied is shown. The transition probability associated with this is *T*(*v*′|*v*) =(1-*ρ*_*x*_)/(3-*ρ*_*x*_).

The transition probability for a myopic agent is

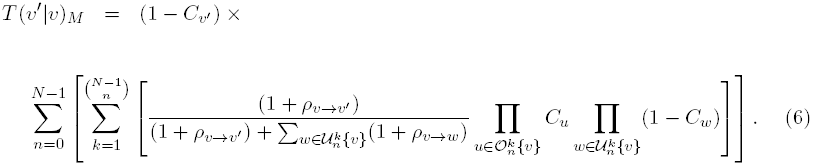

The subscript *M* denotes that this is the transition rate for myopic agent interaction.

### 2.3 Population-level density continuum equations

We have described a range of transition probabilities that represent known cell-cell interactions. In this section we derive corresponding continuum approximations for the spatial and temporal evolution of the population density and expected displacement of a tagged agent on both static and growing domains, and then compare our approximations with the results of our discrete model simulations.

#### 2.3.1 Static domains

We begin by deriving a continuum approximation for a static domain with agents that undergo blind interactions [35]. To derive our continuum approximation we proceed as follows. If the change in the average occupancy of site *v* during the time interval [*t*, *t*+ *τ*) is denoted by *λ*〈*C*_*v*_(*t*)〉 and the change in average occupancy due to a transition from *v*→*v′* is *T*(*v′*|*v*) 〈*C*_*v*_〉 [35], then by summing over all members of the nearest neighbourhood set 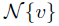 the discrete conservation equation for agents with blind agent interactions can be written as

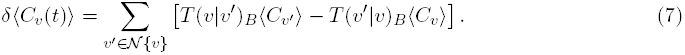

In Eq. (7) our transition rate, *T*(*v*|*v′*)_*B*_, is now treated as an ensemble average transition rate for blind agent interactions. That is, Eq. (7) gives the expected change in occupancy over many realisations, assuming independence of lattice site occupancy. To derive a partial differential equation (PDE) describing the spatio-temporal evolution of the agent population density from Eq. (7) we make a change of variables, such that *v* → (*x*, *y*) and 〈*C*_*v*_(*t*)〉 → *C*(*x*, *y*, *t*), and then perform Taylor series expansions on Eq. (7). It has previously been shown [21, 41, 42] that Eq. (7) is related to a PDE in the limit as Δ → 0 and τ → 0, such that Δ^2^/τ is held constant. Thus, the spatial and temporal evolution of the continuous density, *C*(*x*, *y*, *t*), for a population with blind agent interactions on a two-dimensional square lattice is governed by

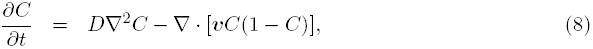

where ***v*** =(*v*_*x*_,*v*_*y*_) is the advective velocity with components

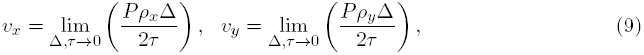

and diffusivity given by

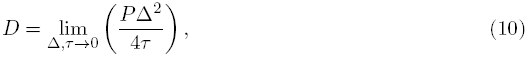

which implies *ρ*_*x*_ = *O*(Δ) and *ρ*_*y*_ = *O*(Δ) [40].

If we follow the same procedure for the adhesive, repulsive and myopic agent interaction transition probabilities, that is Eqs. (3) and (6), we can derive equations for the spatial and temporal evolutions of the density, *C*(*x, y, t*), for adhesive, repulsive and myopic interactions. Further details are given in Appendix A. These equations are listed in Table 1.

**Table 1:** Population density equations for adhesive, repulsive and myopic agent interactions.

| Agent interaction | Population density equation |
| --- | --- |
| Adhesion ($\alpha > 0$) and repulsion ($\alpha < 0$) | $\begin{aligned} \frac{\partial C}{\partial t} = & D \nabla \cdot [(1 - 8\alpha C + (5\alpha + 13\alpha^2)C^2 - (10\alpha^2 + 6\alpha)^3 C^3 + 5\alpha^3 C^4) \nabla C] \\ & - \nabla \cdot \left[ v C \left( \frac{1}{5}(5\alpha^3 - 2\alpha^2)C^4 - \frac{1}{4}(4\alpha^3 + 10\alpha^2 - 4\alpha)C^3 \right. \right. \\ & \left. \left. + \frac{1}{3}(9\alpha^2 + 5\alpha - 2)C^2 - 3\alpha C + 1 \right) \right]. \end{aligned} \tag{11}$ |
| Myopic | $\begin{aligned} \frac{\partial C}{\partial t} = & D \nabla \cdot \left[ \left( 1 + \frac{8}{3}(C + C^2 + C^3) - 5C^4 \right) \nabla C \right] \\ & - \nabla \cdot \left[ v C \left( \frac{1}{9}(13C^4 - 12C^3 - 6C^2 - 4C) + 1 \right) \right]. \end{aligned} \tag{12}$ |

#### 2.3.2 Growing domains

We now include domain growth, as illustrated in Fig. 1, into the continuum approximations of our discrete models. To do so we define the two adjacent column sites of *v*, i.e. (*i, j* − 1) and (*i, j* + 1) as *v*_*L*_ and *v*_*R*_, respectively. The two adjacent row sites of *v*, i.e. (*i* − 1, *j*) and (*i* + 1, *j*) are defined as *v*_*U*_ and v_*D*_, respectively. Ignoring agent movement, we can write a difference equation to account for the effect of growth in the *x* direction on the average occupancy status of site *v*, that is

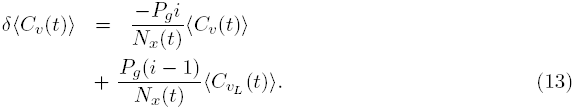

Here, *P*_*g*_ represents the probability of a growth event in the time duration [*t*, *t* + *τ*). The terms on the right-hand side (RHS) of Eq. (13) represent the ways in which a growth event can affect the occupancy status of site *v*. The first term on the RHS of Eq. (13) represents site *v* being occupied and a growth event occurring to the left of, and including, site *v*. The second term on the RHS of Eq. (13) represents a growth event occurring to the left of site *v* and causing site *v* to become/remain occupied. If we transform Eq. (13) from discrete to continuous variables and perform Taylor series expansions, noting that

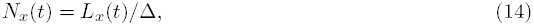

and

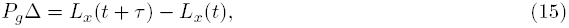

we obtain

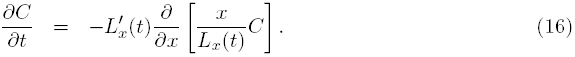

Here, 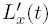 represents the growth rate of our domain in the positive *x* direction. This is the same equation derived by Hywood et al. [29]. Importantly, in the derivation of Eq. (16) we do not explicitly take Δ, *τ* → 0. This approach is reasonable as Δ is non-zero in the discrete simulations. This growth term can now be added to the previous equations describing the evolution of population density, Eqs. (8), (11) and (12), to represent the effects of domain growth. For instance, with the addition of growth Eq. (8) becomes

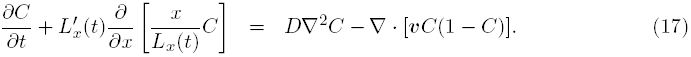

### 2.4 Cell trajectories

Initially, we derive equations for the expected displacement of a tagged agent on a two-dimensional square lattice on a static domain.

#### 2.4.1 Static domains

The expected displacement of a blind tagged agent at site *v* in the positive *x* direction given its position along the *x* axis is *p*_*x*_ at time *t* during the time-interval [*t, t* | τ) is [21]

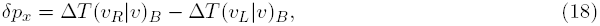

and in the positive *y* direction, given its position in the *y* axis is *p*_*y*_ at time *t*, is

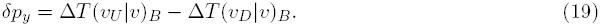

That is, the site length,Δ, multiplied by the transition probability, *T*(*v′*|*v*)_*B*_. It is important to mention here that a simplifying assumption has been made whereby the expected displacement is written solely in terms of an agent’s expected position at time *t*. If we rearrange and divide by *τ*, with Δ^2^/τ held constant as Δ,τ → 0 simultaneously, it has been shown by Simpson et al. [21] that for an agent with blind agent interactions

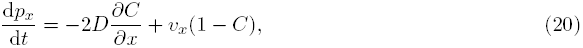

and

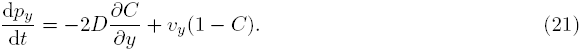

where *C*, *D*, *v*_*x*_ and *v*_*y*_ are defined as previously. If we follow this same procedure with the relevant transition probabilities, Eqs. (3) and (6), we can the derive equations for the expected displacement of agents with adhesive, repulsive and myopic cell interactions. The resulting equations for the expected displacement of a tagged agent are listed in Table 2, including the analogous equations for displacement in the *y* direction. Solving Eqs. (20)-(25) gives the expected displacement of a tagged agent initially at [*p*_*x*_(0),*p*_*y*_(0)]. As can be seen from Eqs. (20)-(25) the rate of change in expected displacement depends explicitly on the average agent density. It should also be noted that if *α* = 0 in Eqs. (22) and (23) we recapitulate expected displacement equation for a blind agent.

**Table 2:** Expected displacement of a tagged agent initially at [*p*_*x*_(0),*p*_*y*_(0)] for adhesive, repulsive and myopic agent interactions.

| Cell interaction | Expected displacement equation in the $x$ direction |
| --- | --- |
| Adhesion/repulsion | $\frac{dp_x}{dt} = -2D(1-\alpha)(\alpha C - 1)^2 \frac{\partial C}{\partial x} + v_x(C - 1)(\alpha C - 1)^3. \quad (22)$ $\frac{dp_y}{dt} = -2D(1-\alpha)(\alpha C - 1)^2 \frac{\partial C}{\partial y} + v_y(C - 1)(\alpha C - 1)^3. \quad (23)$ |
| Myopic | $\frac{dp_x}{dt} = -2D \left( \frac{4}{3} \right) (1 + C + C^2) \frac{\partial C}{\partial x} + \frac{v_x}{9} (C - 1)^2 (9 + 14C + 13C^2). \quad (24)$ $\frac{dp_y}{dt} = -2D \left( \frac{4}{3} \right) (1 + C + C^2) \frac{\partial C}{\partial y} + \frac{v_y}{9} (C - 1)^2 (9 + 14C + 13C^2). \quad (25)$ |

#### 2.4.2 Growing domains

To add growth to Eqs. (20)-(25) we proceed in the following manner. Ignoring agent movement, the expected displacement of an agent at position *p*_*x*_(*t*), at time *t*, in the positive horizontal direction during the time interval [*t*, *t* + τ) due to growth is

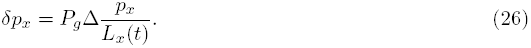

On dividing and taking Δ and τ → 0, Eq. (26) becomes

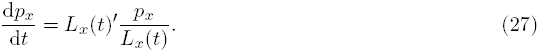

As with the population density equations we simply add this growth term to the expected displacement equations previously derived. For instance, with domain growth Eq. (20) becomes

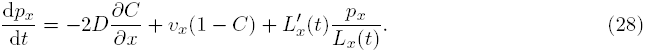

## 3 Comparing continuum approximations to discrete model results

We now compare our continuum approximations with the results of our discrete models. The results presented here are not exhaustive, however they serve to present our key findings. For static domains simulations are performed on a square lattice with *L*_*x*_*(t)* = 200 and *L*_*y*_ *(t)* = 20, with Δ = 1. For growing domains one site is added to each row per time step for all simulations (i.e. *P*_*g*_ = 1). In the case of P_g_ < 1, for instance P_g_ = 0.5, one site would be added to each row every two time steps. All sites between 85 ≤ *x* ≤ 115 are initially occupied with an agent, and an agent at site (114, 10) is tagged. We chose this initial location for our tagged agent as it emphasises the differences between different types of agent interaction. All simulations have periodic boundary conditions at *y* = 0 and *y* = *L*_*y*_ (*t*), and no-flux boundary conditions at *x* = 0 and *x* = *L*_*x*_(*t*). Together these conditions allow us to treat the system as translationally invariant in the *y* axis, hence we can write 〈*C*(*x*,*y*,*t*)〉 = 〈*C*(*x*,*t*)〉.

In all simulations we consider three values of bias parameter *ρ*_*x*_ = 0, 0.1, 0.5, with *ρ*_*y*_ = 0 for all simulations. The density profiles from the discrete models, 〈*C*(*x*, *t*)〉, are obtained by averaging the column occupancy for *R* identically prepared realisations of the discrete model. That is,

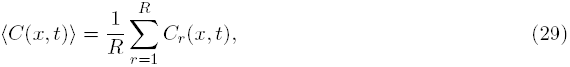

where *C*_*r*_(*x*, *t*) represents the averaged column occupancy in a single simulation at time *t*. We also we calculate the average displacement of the tagged agent over *R* identically prepared realisations of the discrete simulation, that is

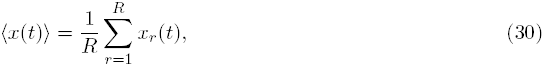

where *x*_*r*_(*t*) represents the displacement of a tagged agent in simulation *r* at time *t*. Comparisons of simulation and continuum results are presented at *t* = 100 (*R* = 100) for static domains, and *t* = 200 (*R* = 500) for growing domains. We generate additional realisations on the growing domain as we simulate the discrete model for a longer time, and the growth mechanism adds additional stochasticity to the discrete model simulation results.

Finally, we introduce a measure for the ‘concavity’ of the expected displacement profile of our tagged agents. To do so we draw a straight line between the initial and final position of a tagged agent, and subtract the area beneath the straight line from the area beneath the expected displacement trajectories of the agent (using the trapezium rule). This difference is then normalised with respect to the distance the agent has travelled. The concavity measure is illustrated graphically in Fig. 4. We will show that with this measure it is possible to identify different types of agent interaction.

**Figure 4:**
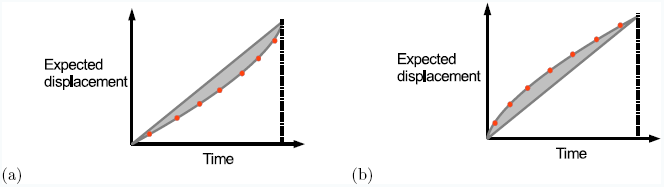
Concavity measure. The dotted line is the trajectory of a tagged agent. A straight line has been drawn connecting the start and final point of the tagged agent. The region between these two lines is shaded and used to calculate the concavity measure. This difference is then normalised with respect to the distance the agent has travelled. (a) Negative concavity, (b) positive concavity.

On a static domain the population-level continuum Eqs. (8), (11) and (12) are solved with a centred-in-space finite difference method and implicit Euler method with Picard iteration with *δt* = 0.05 and *δx* = 0.5 for time integration, and absolute error tolerance ε= 10^−8^. From the solutions of these equations we approximate *∂C/∂x, ∂C/∂y* and *C* using linear interpolation, and then solve Eqs. (20), (22) and (24) with initial position [*p*_*x*_(0), *p*_*y*_(0)] as in [21].

For simulations on a growing domain implement the change of variables used by Crampin et al. [44], that is ξ = *x*/*L*(*t*) and *t* = *τ*. This allows Eqs. (8), (11) and (12) to be solved on a fixed domain [32, 44]. Growing domain equations are solved with a centred-in-space finite difference method and implicit Euler method with Picard iteration with *δτ* = 0.05 and *δξ* = 0.1 for time integration, and absolute error tolerance ε= 10^−8^.

### 3.1 Static domain

We begin by testing the accuracy of our continuum approximations of our discrete model on static domains.

#### Adhesive agent interactions

Fig. 5 row (a)-(b) shows results from discrete model simulations and the corresponding continuum approximation for blind agent interactions. These results have been presented before [21], however, we briefly describe them here. The averaged density profile of the discrete model simulations, taken from 100 repeats, compares well with the solution of Eq. (8) for all values of *ρ*_*x*_, as can be seen in Fig. 5 row (a). The effect of the non-linear advection term introduced into Eq. (8) by a non-zero *ρ*_*x*_ is seen in the asymmetry of the density plots, whereby the agent density skews to the right and reduces in height more quickly compared to agents without motility bias. Fig. 5 row (b) shows that the expected displacement of the tagged agent predicted by the solution of Eq. (20) compares well with the discrete simulation for all values of *ρ*_*x*_. Importantly, the expected displacement depends on the initial position of the tagged agent [21]. We studied different initial tagged agent positions, and found that expected displacement characteristics were conserved for all interactions, although reduced in intensity, as the initial position of the tagged agent approached the centre of the initial condition i.e. positions (100-101,1:20) on the lattice [21].

**Figure 5:**
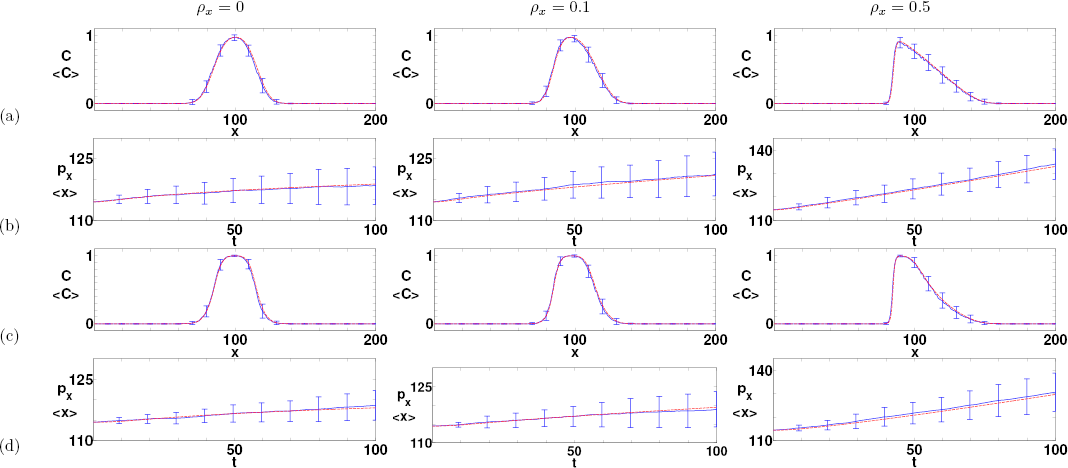
(Colour online). Agent population density and expected displacement for agents with blind and adhesive interactions at *t* = 100. Row (a) shows the averaged density profiles, 〈*C*〉 (solid blue), of the discrete model simulation in comparison with the solution of Eq. (8), 〈*C*〉(red dashed), for blind agent, interactions. Row (b) shows the expected displacement of the tagged agent, for blind interactions. Discrete model simulation results (blue solid) are compared with continuum Eq. (20) (red dashed). Row (c) shows the averaged density profiles, 〈*C*〉 (blue solid), of the discrete model simulation in comparison with the solution of Eq. (11), 〈*C*〉 (red dashed), for adhesive interactions (*α* = −0.2). Row (d) shows the expected displacement, for the tagged agent, for adhesive interactions. Simulation results (solid blue) are compared with continuum Eq. (22) (red dashed). All simulations are averaged over 100 Monte Carlo realisations with Δ = τ =1. All continuum models are solved numerically as described in the text, and error bars display the standard error of the mean of the discrete model simulation.

Fig. 5 row (c) shows a comparison of the discrete model simulation and continuum approximation solution for agents with adhesive interactions with *α* = -0.2. Like the blind case, the averaged density profiles of the discrete model simulations and the solution of Eq. (11) compare excellently for all values of ρ_*x*_ for this value of *α*. However, compared to agents with blind interactions, agents with adhesive interactions spread less, due to fewer movements being attempted. Importantly, the density profile for the agent adhesion process is convex on its RHS in the presence of motility bias (the direction of positive bias). This is due to two reasons: (i) agents with adhesive interactions have dispersed less by *t* = 100; (ii) the adhesive interaction means once an agent has left a high density region it is more likely to move again, resulting in the density skewing in the direction of positive motility bias. Finally, Fig. 5 row (d) shows that the evolution of the expected displacement for the tagged agent predicted by Eq. (22) compares well with the simulation data for all values of ρ_*x*_. In the case of agent adhesion the expected displacement of the agent is less than for an agent with blind interactions, as would be expected.

#### Repulsive agent interactions

We now study our repulsive-type agent interactions, that is, agents with repulsive and myopic interactions. Fig. 6 row (a) shows the averaged density profile for the discrete model simulation and continuum approximation solution for agents with repulsive interactions for *α* = 0.2. Compared to agents with blind interactions, agents with repulsive interactions spread more due to movement probability being ‘redistributed’ from occupied sites to unoccupied sites in the nearest neighbourhood set. As with blind and adhesive interactions the density profiles of the averaged simulation data and solution of Eq. (11) compare excellently for all values of *ρ*_*x*_ for *α* = 0.2. This excellent agreement between the continuum approximation and discrete model is maintained for any 0 < *α* < 0.2. When this threshold is passed the condition stipulated by Eq. (4) is violated and the continuum approximation becomes inaccurate (dependent on the value the motility bias, *ρ*_*x*_).

**Figure 6:**
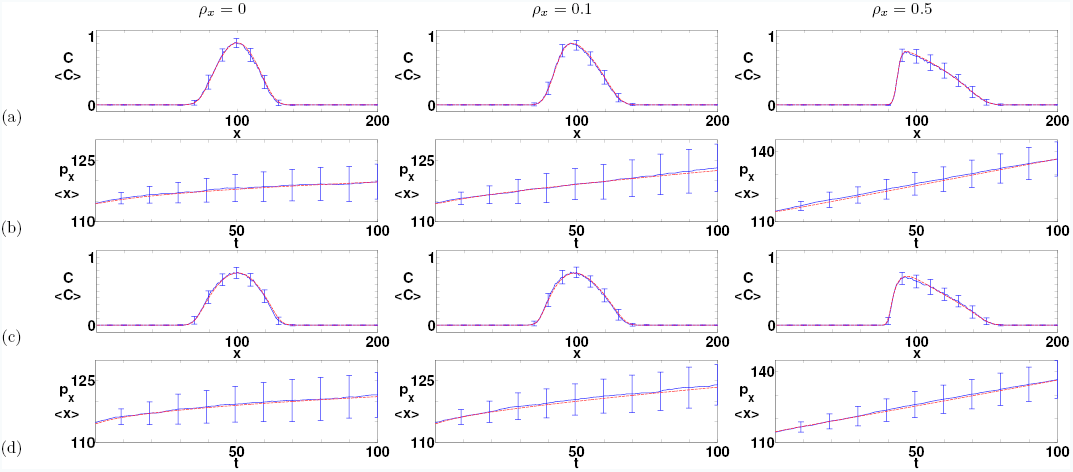
(Colour online). Agent population density and expected displacement for agents with repulsive and myopic interactions at *t =* 100. Parameters and figure descriptions are the same as those in Fig. 5. Rows (a)-(b) are repulsive agent interactions *(*α* =* 0.2) and Eq. (11) and (22), respectively. Rows (c)-(d) are myopic agent interactions and Eq. (12) and (24), respectively.

In contrast to agents with adhesive interactions, the density profile for agents with repulsive interactions is concave on the RHS of the agent population in the presence of bias. This is because agents with repulsive interactions have dispersed more by *t* = 100. This is an important observation as visualising the profile of the cells at the edge of a cell population is experimentally tractable. We suggest that looking at the shape of the leading edge could aid researchers in identifying cell behaviours and interactions. For instance, a chemoattractant could be placed in the vicinity of a cell population *in vitro*, and the density profile at the edge of this population could then be measured to test whether there are adhesive or repulsive interactions between the cells. Fig. 6 row (b) shows that the evolution of the expected displacement for the tagged agent predicted by Eq. (22) compares well with the simulation data for all values of *ρ*_*x*_. As predicted the expected displacement of an agent with repulsive interactions is greater than would be expected for an agent with blind or adhesive interactions. Therefore, the magnitude of the displacement of individual agents is indicative of the type of interactions between agents.

Finally, Fig. 6 row (c) shows the discrete model simulations and continuum approximation solutions for agents with myopic interactions. Like all other agent interactions the density profiles of the averaged simulation data and the solution of Eq. (12) compare excellently for all values of *ρ*_*x.*_ As with agents with repulsive interactions the RHS of the agent population exhibits a concave profile. The myopic agent interaction can be thought of in some sense to be a ‘maximised’ repulsive interaction. The expected displacement Eq. (24) also agrees well with the simulation data for all values of *ρ*_*x*_, as can be seen from Fig. 6 row (d). A tagged agent with myopic interactions exhibits the greatest expected displacement by *t* = 100 compared to the other interactions, which we would expect.

In Fig. 7 the evolution of the standard error of the mean (SEM) for the displacement of the tagged agent in the discrete model simulation for all agent interactions has been plotted against time with and without motility bias. It can be seen that without bias the repulsive-type interactions exhibit a larger SEM in the displacement of the tagged agent at all time points in the discrete model simulations, when compared to blind and adhesive interactions. Therefore, we could expect cells with repulsive-type cell-cell interactions to exhibit greater variance in their displacement in cell trajectory experiments. In the presence of motility bias, however, the SEM of different agent interactions becomes less distinguishable. This is the opposite to what is found when considering the population density, in which case it is in the presence of motility bias that different agent interactions become distinguishable. This difference between the collective and individual migration of agents is an interesting and important observation.

**Figure 7:**
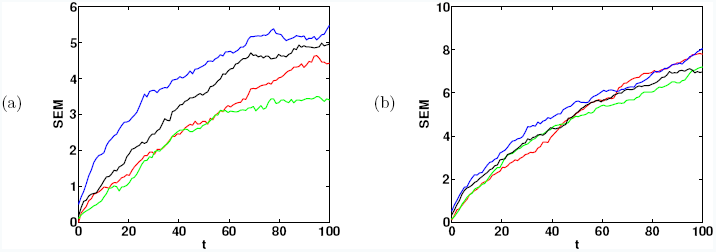
(Colour online). The evolution of the SEM for the position of the tagged agent for all agent interactions. The SEM was calculated from 100 repeats of the discrete model simulation for each agent interaction. (a)*ρ*_*x*_ = 0, (b) *ρ*_*x*_ = 0.5. Blind (red), adhesive (green), repulsive (black), myopic (blue).

### 3.2 Growing domain

We now compare the solutions of our continuum approximations to our discrete model simulations on a growing domain. Fig. 8 row (a) shows a comparison of the discrete simulation and continuum results for agents with adhesive interactions. The density profiles of the averaged simulation data and the solution of Eq. (11) are not as accurate as compared to a static domain for adhesive interactions (although the inaccuracy is only minor). This is noticeable on the left-hand side (LHS) of the agent population for *ρ*_*x*_ = 0.5. However, the agent population retains its convex character in the presence of motility bias on its RHS. This is an important observation, in that the population density for agents with adhesive interactions maintain their population density characteristics in the presence of domain growth. Fig. 8 row (b) shows that the expected displacement of the tagged agent predicted by the solution of Eq. (22) compares well with the simulation data for all values of *ρ*_*x*_. Compared with a static domain the displacement of the tagged agent is much greater, as we would expect due to domain growth.

**Figure 8:**
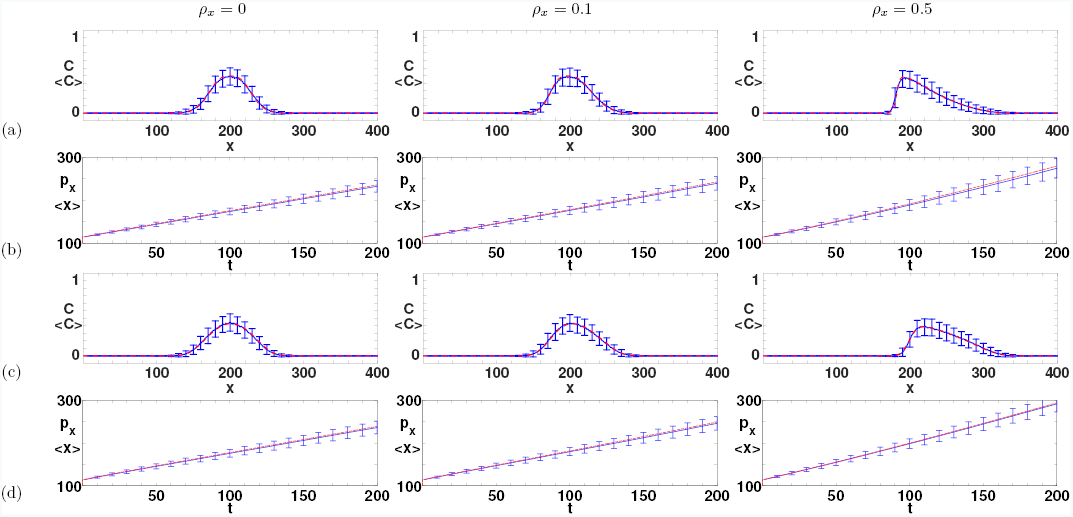
(Colour online). Agent density and expected displacement for agents with adhesive and repulsive interactions on a growing domain at *t =* 200. Row (a) shows the averaged density profiles, ⠩*C*⠪ (solid blue), of the discrete model simulations in comparison with the solution of Eq (11), *C* (red dashed), for adhesive interactions (*α* =- 0.2). Row (b) shows the expected displacement for the tagged agent for adhesive interactions. Discrete model simulation results (solid blue) are compared with continuum Eq (22) (red). Row (c) shows the averaged density profiles, ⠩*C*⠪ (blue), of the discrete simulations in comparison with the solution of Eq (11), *C* (red dashed), for repulsive interactions (*α* = 0.2). Row (d) shows the expected displacement for the tagged agent for repulsive interactions. Simulation results (blue) are compared with continuum Eq (22). All simulations are averaged over 500 Monte Carlo realisations with Δ = τ= 1. All continuum models are solved numerically as described in the text., and error bars are the SEM of the discrete model simulation.

Fig. 8 row (c) shows the spatial and temporal evolution of the population density for agents with repulsive interactions on a growing domain. The averaged density profile of the discrete model simulations compares excellently with the continuum solution of the growth analogue of Eq. (11) for all values of *ρ*_*x*_. This is different to the adhesion process, where the continuum solution does not accurately predict the averaged density profile on the LHS of the population. Like the non-growing domain the density profile for agents with repulsive interactions is concave on the RHS of the agent population. Therefore, both agents with adhesive and repulsive interactions maintain their population density characteristics caused by motility bias in the presence of growth. This is an important consideration when translating experiments carried out *in vitro* to growing domains. Finally, as before, Fig. 8 row (d) shows that the evolution of the expected displacement of the tagged agent predicted by Eq. (22) compares excellently with the discrete model simulation for all values of *ρ*_*x*_.

### 3.3 Summary

These results, for all agent interactions, have shown that our continuum approximations of our discrete model are accurate for static domains, and for the parameters values presented here could be used as our model instead of our discrete model. This is important as discrete simulations are time-consuming and difficult to complete parameter sweeps with. However, with the addition of domain growth our continuum approximations for the spatial and temporal evolution of the population density for agents with adhesive interactions were less accurate, especially on the LHS of the agent population. We found this to be the case for agents with myopic interactions too, but not for agents with blind interactions, which gave results that were similar in accuracy to agents with repulsive interactions. Finally, our continuum approximations for the expected displacement of a tagged agent were accurate for all four interactions for the parameter values presented here (data not shown for blind and myopic interactions).

### 3.4 Tagged agent trajectories

We now briefly discuss the results of our continuum approximations for the expected displacement of a tagged agent in more detail, and how they could be used to identify different agent interactions. Fig. 9 shows the expected displacement of the tagged agent for all interactions when *ρ_x_*= 0. All four trajectories have inflection points, which in the blind case signifies when the tagged agent passes the inflection point in the agent population density profile. This can be seen most easily by looking at Eq. (20) with *ρ*_x_ = 0. If we take the partial derivative with respect to time we obtain

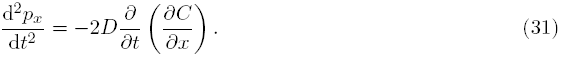

**Figure 9:**
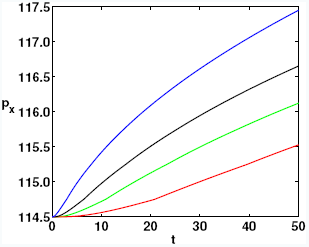
(Colour online). The expected displacement trajectory of a tagged agent in the positive *x* direction for all interactions with respect to time. Eqs. (20), (22) and (24) with *ρ*_*x*_ = 0 and on a static domain. Blind (green), adhesive (red), repulsive (black), myopic (blue).

Additionally, we know that

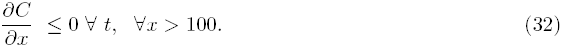

So for d^2^*p*_x_/d*t*^2^ to change sign we require *∂*/*∂t*(*∂C*/*∂x*) to change sign at the position of the tagged agent. This occurs when the tagged agent passes the inflection point in the agent population density profile, as the second derivative at the position of the tagged agent changes in sign. This observation, although qualitative, could be of some use to researchers trying to identify cell properties through cell trajectories. Cells with repulsive cell-cell interactions may be expected to exhibit a convex trajectory in their displacement for a shorter amount of time (black line, Fig. 9), after which they exhibit a concave trajectory. Conversely, the expected displacement trajectory of cells with adhesive cell-cell interactions exhibit a convex shape for a longer amount of time compared to cells with repulsive cell-cell interactions (red line, Fig. 9).

**Figure 10:**
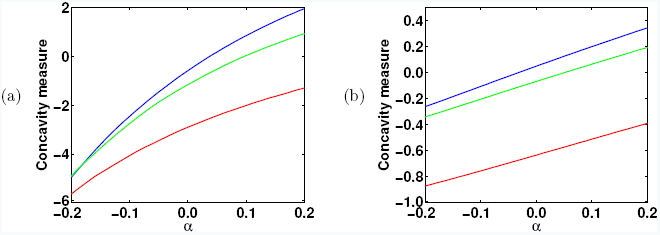
(Colour online) Agent trajectory concavity measure with respect to adhesion parameter *α* for a tagged agent initially situated at (114, 10). (a) Static domain, *ρ*_*x*_ = 0 (blue), *ρ*_*x*_ = 0.1 (green), *ρ*_*x*_ = 0.5 (red). (b) Growing domain, same colour scheme as (a).

Fig. 10 displays how the concavity measure of a tagged agent depends on the adhesion parameter *α*. Fig. 10 (a) shows the concavity measure for a static domain at t = 50. The blue line shows the non-biased case, *∂_x_* = 0. We can see that for most positive values of a (i.e. repulsive interactions) the trajectory has a positive concavity measure. However, as *α* approaches zero, that is, interactions become blind, the concavity measure becomes negative, as for adhesive interactions the tagged agent trajectory is convex. Therefore, we suggest that the concavity measure could be used to identify repulsive and adhesive cell-cell interactions experimentally. In the presence of a strong motility bias, *∂_x_* = 0.5, the concavity measure is negative for all values of *α*. This is important as it suggests that repulsive interactions may not exhibit concave profiles in the presence of motility bias. Meanwhile, Fig. 10 (b) shows the same continuum approximation solutions with the addition of growth at t = 50. Here we see that while growth reduces the magnitude of the concavity measure of the trajectory of a tagged agent with repulsive interactions, it does not make the concavity measure negative. Therefore, trajectory characteristics associated with different agent interactions are conserved in the presence of the growth mechanism implemented in this work.

## 4 Discussion and conclusion

In this work we have extended previously formulated discrete and continuous models for a range of agent interactions, representing known cell-cell interactions in biological systems. These have been implemented on both static and growing domains, using a ‘pushing’ growth mechanism [19, 29, 32]. Our continuum approximations accurately predict the evolution of the averaged population density over a wide range of parameter space. The density profiles resulting from different types of interactions are differently shaped. Agents with myopic and repulsive cell-cell interactions display concave profiles in the presence of a motility bias, whereas agents with adhesive interactions exhibit a convex profile in the presence of motility bias. These profiles were conserved on both static and growing domains, suggesting this behaviour would be conserved in the presence of domain growth, an important consideration when considering whether experimental data collected in *vitro* is applicable to growing domain environments. As discussed in Section 3, visualising the profile of the cells at the edge of a cell population in the presence of a chemoattractant is potentially experimentally tractable, and so this knowledge could help researchers identify different cell-cell interactions.

We have also presented continuum equations describing the expected displacement of a tagged agent with different interaction mechanisms. By studying the expected trajectory of a tagged agent with different agent interactions (Fig. 9) we showed that different agent interactions give rise to different trajectory profiles and magnitudes. For agents with repulsive interactions our tagged agent exhibited a largely concave trajectory compared to an agent from with adhesive interactions. In the presence of strong motility bias the differences in the trajectories of tagged agents with different agent interactions was lost. Moreover, agents with repulsive-type interactions exhibited greater expected displacement compared to agents with adhesive interactions. This is an obvious but important consideration when analysing cell trajectory data, and could be used to identify cell-cell interactions. We also found that repulsive-type agent interactions exhibited a greater SEM when compared to blind or adhesive interactions. Therefore, the SEM could also be used as a potential means by which to distinguish between repulsive-type cell-cell interactions. The difference in the magnitude of the SEM for tagged agents with different agent interactions was lost in the presence of a strong motility bias. This highlights how understanding the effect of different agent interactions in the presence of motility bias is not straightforward. Ironically, the presence of motility bias is what allows us to distinguish between different agent interactions when looking at the population density. However, the presence of motility bias obscures the differences between the trajectories of tagged agents with different agent interactions. This is an interesting difference between analysing population and individual agent level statistics from our discrete and continuum models.

It is important to discuss how the concavity measurement will depend on the time period over which it is measured. This is because the rate of change of the concavity measure reverses in sign in value after an inflection point in the cell trajectory. If we observed a cell with adhesive cell-cell interactions with no directional bias on a static domain at the edge of a cell mass, such as in Fig. 5 row (d), we would eventually expect an inflection point in its expected displacement trajectory (see Fig. 9, red line) [21]. Therefore, it would be necessary to study cell trajectories over an appropriate timescale. The relationship between the variance in cell trajectory data and the timescale of measurement has been previously studied, and it was shown that collecting large numbers of repeats over short timescales is most effective in reducing the variance of cell trajectory data [45]. Therefore, given the nature of cell trajectory experiments we would only recommend use of the concavity measure over short time scales with multiple repeats, as with long time scales both theoretical and experimental difficulties with cell trajectory experiments arise. However, it stands to reason that over short timescales the different concavity properties of cell trajectories with adhesive and repulsive cell-cell interactions presented here could be viable, as long as the experiment is effectively controlled.

A final consideration concerns the uniqueness of possible statistics used to infer cell-cell interactions. That is, could different cell-cell interactions give rise to similar population density and cell trajectory statistics? Although we have not addressed this issue here, it is necessary to be aware of this possibility. That different cell-cell interactions may exhibit similar migratory behaviour is of interest and a topic worthy of further research.

## Acknowledgements

RJHR would like to thank the UK’s Engineering and Physical Sciences Research Council (EPSRC) for funding through a studentship at the Systems Biology programme of the University of Oxford’s Doctoral Training Centre. The authors would also like to thank two anonymous reviewers for their helpful comments.

## Appendix A

To derive Eq. (11) we proceed in the following manner. The discrete conservation equation for agents with adhesive agent interactions can be written as

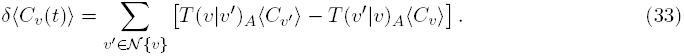

This equation is the same as Eq. (7) in the text, apart from for agents with adhesive interactions as opposed to blind ones. This is indicated by the subscript *A* on the transition probability, *T*(*v*|*v′*)_*A*_. As before, we treat our transition rate, *T*(*v*|*v′*)_*A*_, as an ensemble average transition rate for adhesive agent interactions. That is, the expected transition rate if the simulation is repeated many times. An example two-dimensional cell adhesion transition probability for a movement from (*i*, *j* + 1) → (*i*, *j*) using the ensemble average of many discrete simulations is:

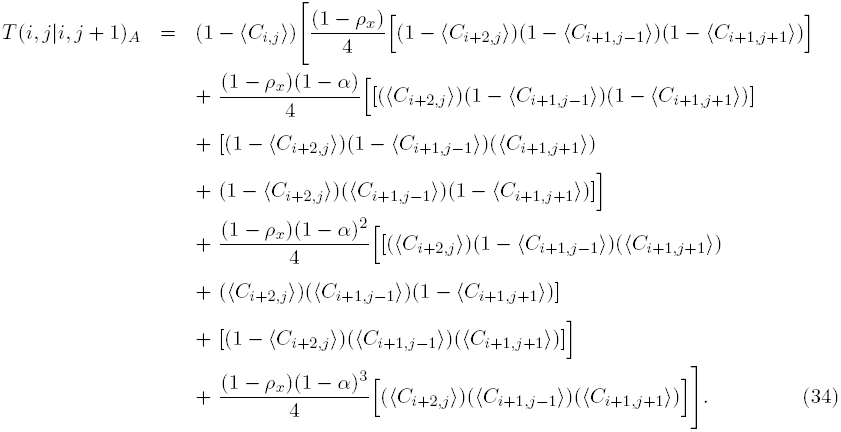

Using Eq. (33), if we perform Taylor series expansions and take the limits as prescribed in the text we arrive at Eq. (11).

